# Dominance of indirect effects of deer populations on soil biodiversity

**DOI:** 10.1101/2025.10.29.685376

**Authors:** Nida Amin, Colin Brock, Aven Beech, Virginia Morera-Pujol, Adam F. Smith, Simone Ciuti, Tancredi Caruso

## Abstract

Deer abundances and distributions have expanded at increasing rates in several regions across the globe due to human-driven land use change, uncontrolled introductions and reintroductions, and insufficient top-down control by human or natural predators. Yet, the impact of unsustainable density of deer on the belowground component of ecosystems is understudied. Ireland is a good model for anthropic environmental context in which the population of large herbivores is not regulated by large predators nor planned hunting management. Through a combination of camera trapping paired with soil and vegetation surveys conducted with high-throughput technology, we monitored 50 sites in Irish woodlands to study the effect of varying sika deer (*Cervus nippon*) relative densities on soil biota. Site-level deer intensity of use correlated with vegetation through an increase in litter and moss quantity and reduction in understory vegetation. Furthermore, the effect covaried with soil pH and cascaded on soil microarthropod, bacterial and fungal communities, with more opportunistic taxa linked to high relative deer densities. We also found that variation in deer intensity of use directly affects vertebrate communities, which shifted towards a dominance of foxes, squirrels, cats and dogs – species more commonly associated to human-dominated landscapes – and a reduction in birds and mustelids, which also cascaded on soil biota. Our results thus support the hypothesis that deer overabundance has a strong potential to drive the structure of soil biota mostly through indirect effects via vertebrate fauna and vegetation, acting as a powerful ecosystem engineer. Future studies will have to experimentally tease apart the mechanisms underpinning the patterns documented in this study.

## 1. Introduction

Over the last two decades ecologists have emphasised the need of linking aboveground and belowground components of terrestrial ecosystems to improve the understanding of their structures, functions, and responses to environmental impacts [1]. In terrestrial ecosystems, plants play the major role in terms of the dynamics of energy and matter, being the primary producers and controlling the flux of nutrients between the atmosphere and the soil [2]. At the same time, large herbivores regulate the structure of vegetation and, with their physical and chemical impact, also affects soil and soil biota [3]. This suggests that changes in large herbivore populations may cascade on and alter the entire ecosystem. For example, excessive deer herbivory has direct negative effects on the diversity and abundance of several plant species, tree seedlings, saplings, woodland herbs, and understory vegetation [4–7] resulting in the prevention of tree regeneration and an increase in the dominance of those plant species that are avoided by deer or are browse-tolerant, which are often invasive [8–11]. Herbivory may also lead to the loss of understory vegetation, which exposes litter and soil surface, increasing wet-drought stress and resulting in soil erosion [12, 13]. At the same time, animal trampling increases soil compaction [5], reduces soil aeration [14], reduces the abundance and diversity of arbuscular mycorrhizal fungi (AMF) with potential loss of AMF beneficial effects for plants under intense grazing conditions [15], and can overall impact the processes of litter decomposition and nutrient cycling [16]. The deposition of animal urine and faecal biomass, together with selective browsing and trampling, can positively affect the composition and structure of aboveground plant species [11].

Herbivory on vegetation by deer overconsumption can also indirectly impact other animals aboveground such as insects, birds, small mammals, reptiles and amphibians [17–21]. This is due to the alteration in availability of food sources, cover from predators and, more generally, the alteration of the micro-habitat conditions and resources that those populations require to remain stable [18, 22]. Other studies have suggested that the effect of deer herbivory on belowground communities is weaker than those on aboveground communities [23]. Yet, long-term herbivory can affect soil food webs directly through waste deposition and trampling, and indirectly through alteration in plant biomass, secondary compounds production in roots and foliage, changes to the soil quality in terms of organic matter composition, thus resulting in a modification of composition and diversity of soil biota and its functions such as nutrient and carbon cycling [24–27].

Given the ecological interactions between large herbivores such as deer, plants and soil, the lack of regulation of herbivores has the potential to modify the structure and function of terrestrial ecosystems. We propose Ireland as a perfect point in case: an increase in deer density in Ireland due to human-driven land use change, and lack of science-based deer management, or other controls such as natural predation, is of particular concern for agriculture and both natural and commercial forestry [28, 29]. This has led to the establishment of fenced areas as a somewhat effective strategy to protect woodland ecosystem from deer-induced damages [30], creating within the same areas with similar ecological conditions a significant disparity in deer relative density where it is possible to disentangle direct and indirect impacts of deer on soil biota.

Here, we focused on woodland dominated by sika deer (*Cervus nippon*), which was introduced to Ireland in 1860 [31]. Since then, deer escapes, releases and translocations have generated “hotspots” with locally high relative density [28, 32] The total deer population in Ireland is unknown, although it is well documented to be expanding and increasing [29, 33]. We selected the major hotspot of sika deer presence (county Wicklow), where, in 2022/23 only, more than ∼19,000 sika deer were legally shot (NPWS official data). We focused on sika deer because they display opportunistic feeding behaviour, enabling them to feed on a range of vegetation [34]. The major component of sika deer diet in Ireland comprises of grass, followed by holly, gorse, heather, bark and saplings [35–37].

To study the effect that sika deer are having on woodland ecosystems, we established a network of 50 sites in the core of the sika deer distribution (obtained from [28] varying in deer relative density, in a gradient from fenced areas with no deer to areas with very high density of deer while keeping a low variance in terms of tree species, tree density, and canopy cover. In each site, we measured a broad range of biota including mammals and birds (with camera traps), soil microbes (high-throughput sequencing) and microarthropods (microscopy), and vegetation structure and composition. Our overarching goal was to collect evidence on the potential of sika deer population to structure aboveground and belowground biodiversity in woodland and quantify the direct and indirect effects of high levels of deer populations on soil biota. We hypothesised that high deer abundance is associated with either direct, and indirect or combined direct and indirect but predictable structural changes in the diversity and composition of soil biota and specifically aimed at collecting evidence not only on these effects but also on the relative important of direct and indirect effects.

## 2. Materials and methods

### 2.1 Survey design and site selection

Our study area was located within a known sika deer hotspot area across Co. Wicklow and northeast Co. Carlow (Fig. 1), in the east of Ireland [28, 29, 32]. Permission to use these sites was granted in advance from both public and private landowners (e.g., property owners, Coillte and National Parks & Wildlife Service). Our selected sites were spread across and beyond areas known to have high deer relative densities based on the most recent and up-to-date estimates of sika deer distribution and relative densities [28, 29]. Sampling stations (n = 50) were chosen to also include fenced forested areas (fenced for at least 2 years prior to the data collection to allow the soil recover from previous deer presence) with similar ecological conditions to the unfenced sites. In total, 6 out of the 50 sites were fenced to exclude deer entering these patches of forest. Note that we used [28] estimates to select our sites across a gradient of predicted relative densities combined with information on fenced areas (i.e. exclusion zones); actual presence and intensity of use of a site by deer was estimated *ex-novo* using the camera traps. We avoided forest areas dominated by single tree species plantations (i.e., Sitka spruce) and selected forests with a native broadleaf composition. The average distance between sites was 1.9 km, the minimum distance between any two sites was 190 meters, and the maximum distance between one site and the next closest site was 7 km. In general, sites were selected at a distance greater than 1.5 km to guarantee spatial independence. Lower distance between sites was allowed only when sampling a fenced site without deer access and a similar site located close by with full deer accessibility.

**Fig. 1.**
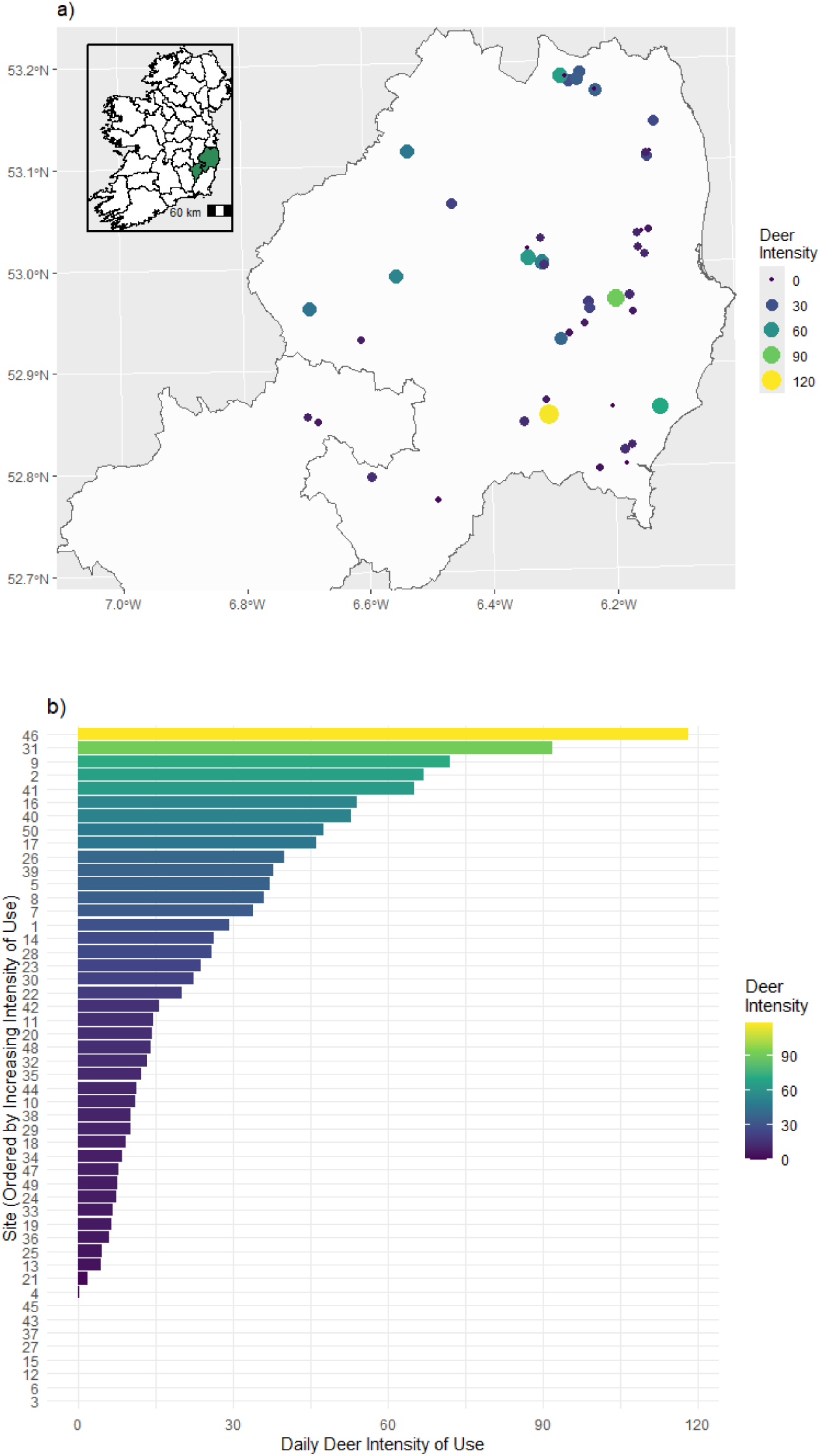
Gradient of daily deer intensity rates across 50 sites in Co. Wicklow and northwest Co. Carlow, Ireland. (a) Shows the spatial distribution within Wicklow and Carlow regions. Each point represents a sampling site with the size and colour highlighting the differences in daily intensity use. Smaller and darker colours represent low daily intensity of use and larger brighter colours represent higher daily intensity of use. (b) Shows a bar plot for each site within the study area. X-axis and colour gradient highlights daily intensity rate of deer at each site, with dark blue colours representing low deer intensity and bright yellow colours representing high deer intensities. Y-axis represents the site numbers, ordered by increasing daily intensity rate to highlight the deer intensity gradient across all 50 sites.

### 2.2 Camera Traps

We deployed Browning Trail Camera Recon Force Elite camera traps in each of our 50 sites. The following settings on the camera traps were configurated in all 50 cameras: photo quality set to medium (8MP), photo delay set to as fast as possible (1 second), multi-shot mode set to 3 shot rapid-fire. All remaining settings were left at the default option. They always monitored continuously and once triggered by a moving object would take three rapid photographs every 1 second for as long as the camera was triggered. Camera traps were deployed at the sampling locations within our fenced and unfenced sites in the spring from the 24th of April to the 4th of May 2023, and retrieved from the 24th of August to the 12th of September 2023. We fixed the cameras onto trees at a height of roughly 50cm parallel to the ground, facing north and without bait.

We created our camera trap dataset by first running a classification of all images using the DeepFaune software (Version 1.0.1), a deep learning model programmed to classify many European species [38]. The taxa selected for classification within DeepFaune included: badger, bird, cat, cow, dog, fox, goat, hedgehog, human, lagomorph, micromammal, mustelid, red deer, sheep, squirrel, vehicle. While sika deer are the most common species in our area, and fallow deer (*Dama dama*) are also found in the study area, neither were included in the version of DeepFaune we used. However, we found that the software would most often predict another deer species regardless, and so we used the more encompassing definition “deer” (i.e., Cervidae) to describe any species observed in an image. Although DeepFaune was extremely useful in a first round of classification of our images, as well as creating the appropriate structure for our dataset, we manually corrected or validated all the images that were classified by DeepFaune to ensure that we had the correct taxa for each image to calculate a site intensity of use metric. We also added observation data where we had groups of deer. Intensity of use for each site was calculated for deer and all other animal species/groups identified through the camera traps detailed above. Intensity of use of a site by deer was calculated by the number of images recorded for a site divided by the number of days monitored, and was used as a proxy for the mechanical use of the site by deer. We also calculated an encounter rate for each site, which is the number of independent observations of a species per survey day, and is a more often utilised proxy for density (e.g., Palencia et al. 2021 and references therein). However, we found that site intensity of use and encounter rates of deer were highly collinear (Supplementary Information, Fig. S1), so we took only intensity of use further because of its higher relevance to our research question (i.e., the intensity of use at the site level expected to be a proxy of the amount of time a single and/or multiple deer would spend grazing, browsing, trampling-walking-running, including waste deposition).

### 2.3 Aboveground Vegetation and Habitat

Aboveground vegetation surveys were conducted in the summer months when the vegetation and canopy cover was full. Quadrat surveys were used to measure understory vegetation. Quadrats were randomly placed in six positions across a 180° angle in front of the camera and percentage estimates for the understory functional groups were estimated within the boundaries of the quadrat. Understory functional groups included: grass, forbes, legumes, moss, ivy, litter, shrubs, ferns seedlings and mushrooms. Percentage estimates were later averaged for each functional group per site.

Three concentric circles were made with the centre point 3m in front of the camera in the northwards direction of where the camera was facing (ForstBW, 2012). Each circle had a radius of 1.5m, 2m, and 6m respectively. This method of concentric circles captures different stages of tree growth depending on their diameter at breast height (DBH) or tree height within varying sized circle plots. Tree regeneration was measured in circle 1 and was defined by any tree with a DBH <10cm and height <1.3m. Tree regrowth was measured in circle 2 and was defined by any tree with a DBH <10CM and Height >1.3m. Finally, tree stock was measured in circle 3 and was defined by any tree with a DBH >10cm. Densities per hectare were later calculated with equations detailed in (ForstBW, 2012). Images were taken with a camera facing perpendicular to the ground at the centre point to measure canopy cover percentages. The images were analysed in R 4.3.2 (R Core Team 2023) using the “EBImage” package (Pau et al., 2010).

### 2.4 Soil sampling

Soil samples were collected from the same quadrats used for vegetation and then pooled to form a composite sample representing the site. More in detail, given the scale of the deer impact and the need of replication at large scale, we did not invest in sampling for intra-site variation (a few meter scale) in soil biotic and properties ecology but rather on inter-site variation. Each individual sample (six samples per site, pooled to form the site composite sample) was collected with a 10cm deep by 2.5cm diameter Eijkelkamp gouge auger. On the same quadrat, 6 smaller cores (10cm depth by 1.3cm diameter Eijkelkamp gouge auger) were also collected and combined into a composite sample for the site. This sample was kept at −20°C for subsequent molecular and chemical analysis. The soil microarthropods samples were processed the same day on Burlese-Tullgren funnels for one week, which was sufficient to completely dry the sample. Animals were collected in 75% ethanol. Soil organic matter was determined by loss on ignition: air dried soil samples were ignited in a muffle furnace at 500°C for a minimum of 5 hours, and until the sample ceased to decrease in weight.

### 2.5 Soil biota

Microarthropods were sorted by hand under the microscope at a Leica M165 C fitted with the Leica MC170 HD camera attachment using a Leica 10450028 Planapo 1.0X lens at magnifications ranging from 1.25X to 12X depending on the size of the animal or particulate being identified. Animals were initially sorted into broad (mostly subclasses and orders) but functionally meaningful taxonomic categories such as collembola, oribatid mites, predaceous mesostigmata mites, and other high level (mostly Order) groups of microarthropods such as millipedes and centipedes or insect larvae (identified at the order level). All mites (a minority) which did not fit within the oribatids, or predaceous mesostigmata were classified as ‘other mites’. These mites were dominated by Prostigmata and some Astigmata; despite the latter are now established as a lineage of oribatids [39, 40], given their little abundance in our soils and specific ecology, we kept them separate as done in many studies in the past. Enchytraeids, earthworms and nematodes were occasionally also found and recorded in the samples, however given the extraction methods target microarthropods, they were excluded from the data analysis.

For PLFA analysis of bacteria and fungi, soil was sieved and homogenised through 2mm mesh and, after drying, 5g used to extract lipids according to [41], lipids were fractioned into neutral lipids, glycolipids, and polar lipids on a silica acid column (Bond Elut, Varian Inc., Palo Alto, CA, USA). To do that, we applied a series of elution with chloroform, acetone and methanol. The PLFAs are contained in the methanol fraction, which was subjected to mild alkaline methanolysis to obtain free fatty acid methyl ester from the PLFAs, followed by identification and quantification on a gas chromatograph. We used the sum of the PLFAs i15:0, a15:0, i16:0, 10Me16:0, i17:0, a17:0, cy17:0, 10Me17:0, 10Me18:0 and cy19:0 as an indicator of bacterial biomass while the PLFA 18:2ω6 was used as an indicator of saprotrophic fungi [41–43].

To profile bacterial and fungal communities, DNA was isolated from 250mg of soil sample using manufacturer’s protocol of the DNeasy PowerSoil Pro Kit (QIAGEN, USA). The quality and concentration of extracted DNA was measured using BioDrop (Labplan Ltd.). The DNA samples were diluted to obtain 10ng DNA/μl of distilled H2O. The primer pair with overhang Illumina adapters 341F/806R were used for PCR amplification of V3-V4 region of bacterial 16S rRNA gene [44]. For amplification of ITS2 region of rRNA cistron the primers 5.8fun/ITS4fun with overhang Illumina adapters were used [45]. The PCR mixture (30 µl) comprised of Thermo Scientific DreamTaq™ Green PCR Master mix (2X), primers (10 µM), Mgcl2 (25nM), Bovine Serum Albumin (BSA - 10mg/ml), Dimethyl Sulfoxide (DMSO - 5%), 3 µl of DNA template (10ng/µl) and 7.95µl of distilled water. For 16S amplification, the PCR cycle comprised of a 10min initial denaturation step at 95°C, followed by 20 cycles including 40s denaturation at 95°C, 30s annealing at 60°C, 1min extension at 72°C and 10min of final extension step at 72°C. For ITS amplification, the same PCR cycle was used except annealing step that was done at 57°C for 30s. The 1 µl of PCR product was used in a second PCR step. The second PCR was performed using similar thermocycling conditions and consists of 15 PCR cycles. The reverse primers comprised of additional sequences to integrate index primers and Illumina multiplexing sequence. The final PCR products were purified, quantified, standardised and sequenced using 2×300 bp paired-end Illumina MiSeq sequencer.

### 2.6 Bioinformatics analysis of sequencing data

The RAW 16S and ITS data was processed for bioinformatics analysis using QIIME 2 workflow (2024.2) [46]. The RAW demultiplexed paired-end sequences were imported in Qiime2 environment using qiime tools import and Casava 1.8 demultiplexed (paired-end) format. For 16S data, the primers were removed with q2-cutadapt plugin and trim-paired command that employs cutadapt (v4.6 with Python 3.8.15) within QIIME 2 environment [47]. ITS primers were removed using Q2-ITSxpress plugin (v1.8.1) and trim-paired command [48]. For trimmed ITS sequences, no further quality filtration was applied after ITSxpress. However, the trimmed 16S sequences were filtered to remove bases with average quality score <35. The paired-end reads were joined at mean length (417 ± 14.8 bp, 16S) and (240 ± 18.22 bp, ITS), non-overlapping regions and chimeras were discarded, sequences were clustered into amplicon sequence variants (ASVs) using q2-dada2 plugin and denoise-paired command [49]. The taxonomic assignment was based on the latest release (138) of SILVA reference database for 16S rRNA representative sequences and for ITS, latest release (ver10) of UNITE ITS database was used with feature-classifier plugin and classify-sklearn method in QIIME2 [50, 51]. The resulting artifacts of DADA2 step including FeatureTable [Frequency] and Feature-Data [Sequence] were further filtered to remove non-bacterial and unassigned sequences, and those assigned only at phylum-level using QIIME 2 taxonomy-based filtering step using filter-table command. The singletons were discarded based on Identifier-based filtering step using filter-features and filter-seqs commands. A total of 28 reads (ITS) and 743 reads (16S) were observed in the negative control, which were removed from further analysis. For 16S dataset, in total 6,545,431 reads, 132,334 average read per sample and 12,250 unique bacterial ASVs were observed. For ITS dataset, 4,784,681 reads, 93,931 average read per sample and 3,642 unique fungal ASVs were observed in 50 samples.

### 2.7 Statistical analysis

The dataset included a large number of variables measured to describe multiple components of the ecosystem, including the vegetation (canopy, tree richness, DBH avg, total density, reg density (ha), grass, forbes, moss, ivy, litter and seedlings), soil properties (acidity and OM), vertebrate fauna (badger, bird, cat, dog, fox, hedgehog, lagomorph, micromammal, mustelid and squirrel encounter rates from camera traps), soil microarthropod fauna (24 groups), the diversity, composition and taxa relative abundance of bacteria and fungi (high through-put sequencing data). The number of sites was relatively limited compared to the number of variables measured in the study and the key goal of the study was to demonstrate an association between deer intensity of use and soil biodiversity at the ecosystem level. We therefore adopted a multivariate analysis strategy to combine variables of the same type, and highly correlated, into indicators of the major components of the system under-investigation. Specifically, the three major groups of response variables were the vertebrate fauna (mammals and birds), the soil microarthropods (soil fauna), and the microbes. The two key drivers of these three sets of variables were, by construction of the field survey, the gradient in deer intensity and vegetation, which varied between sites in relation to a number of factors difficult to fully control such as the age of the woodland, local management, and also the potential long-term impacts of deer itself on each site vegetation.

The ASV bacterial and fungal data were rarefied to the lowest number of reads in the data (23,045 reads for 16S and 53,966 reads for ITS) to avoid sampling biases due to uneven sequencing depth. The bacterial and fungal ASVs count data, with the sample metadata and taxonomies, were converted to phyloseq-formats using *phyloseq* package in R [52]. Changes in the composition were quantified with the Bray–Curtis dissimilarity matrix on standardised and rarefied ASVs count data and visualised using Principal coordinate analysis plot. The impact of deer on bacterial and fungal community structure was tested with permutational Analysis of Variance (Adonis2). To identify taxa responding to deer, the *phyloseq_to_deseq2* function of *DESeq2* package [53] was used to convert phyloseq-format datasets into DESeqDataSets, which enabled the calculation of dispersions estimates between deer presence/absence. The distribution used in the linear model was the negative binomial and differences were visualised using the logarithmic fold change (log2FoldChange) to compare each ASV abundance in the presence or absence of deer (DESeq function). The Wald test with Benjamini-Hochberg correction (*P* < 0.05) was used to identify responsive taxa. We focused on the differential ASVs that displayed a log2FoldChange >0.5 or <-0.5 at the adjusted *P* < 0.05, and visualised results using volcano plots with *EnhancedVolcano* package in R [54]. The resulting set of ASVs for bacteria (39 genus-level taxa) and fungi (53 genus-level taxa), and the data containing soil microarthropod fauna (24 species), vertebrates (10 species / species group), and vegetation (11 parameters) were then used for a Principal Coordinate analysis (PCoA) to quantify major gradients of covariation in these sets of variables.

Prior to the ordination, the soil parameters (pH and OM) and vegetation data were standardize to a z-score while the community data (vertebrate fauna, soil fauna, bacteria and fungi) were transformed using Hellinger transformation [55]. Principal Coordinate Analysis was then applied to each individual data. PCoA was conducted using *rda* function of *vegan* package in R. The first PCoA axis labelled as PC1 was used as a descriptor of the groups of variables (bacteria, fungi, soil fauna, vertebrate fauna and vegetation) in further analyses as described below.

We used two main strategies to investigate the relationship between the various sets of variables: Redundancy analysis (RDA) and Structural equation modelling (SEM). RDA was carried out to describe the correlation between each set of variables and some of the variables expected to be mostly related/driving for that set of variables. RDA analysis was done in R using *rda* function of *vegan* package [56, 57]. The adjusted R^2^ (corrected for explanatory variables number) was used to describe the descriptive power of the model. The ordiR2step function with forward selection and 1000 permutations was used to select statistically important variables based on adjusted R^2^ and *P* values [57]. The significance of the RDA model and each variable was tested using anova.cca function. The RDA results were visualised using RDA triplots with Type–I scaling.

Structural equation models were used to evaluate the direct and indirect effects of varying deer densities on the vegetation, soil properties, community structure of vertebrates, and soil microorganisms including fauna, bacteria and fungi. As such, the SEM approach represented the most direct way to address our major goal. We started from the general model topology that assumed a covariance term linking vegetation and deer density, and then tested three possible model structures : Model 1, with direct and indirect effects linking deer to soil properties and biota; Model 2, as model 1 but just with direct effects; Model 3, as model 1 but just indirect effects. See Supplementary Table S1 for further details. SEM was coded and fitted using the *lavaan* R package (version 0.6-17) [58]. Model fit was assessed with the Chi-square (model accepted for *P* > 0.05), the comparative fit index (CFI) ≥ 1.000 and root mean square error of approximation (RMSEA < 0.001). As we used maximum likelihood to fit the models, we could calculate the AIC metric to compare the relative performance of each model structure on the same dataset.

## 3. Results

A total of 226,673 images were collected and classified from the camera traps with deer the most captured animal on the camera traps with over 150,000 (>65%) deer images. Our classification results revealed a gradient of deer intensity of use across our 50 sampling sites (Fig. 1). Importantly, intensity of use was well distributed spatially across different environmental contexts and predicted relative densities (Fig. 1a). Intensity of use by deer ranged from as low as 0.29 images per day of deer to as high as 118.3 images per day (Fig. 1b). Other notable species/animal groups captured were badgers, birds, foxes, micromammals, mustelids and squirrels (Fig. S2, S3). Animals scarcely captured included bats (4 sites), goats (1 site), hedgehogs (3 sites) and lagomorphs (7 sites) with daily intensity of use of 0.6, 4.9, 1.8 and 17.4 (S4). Domestic cats and dogs, as well as humans were captured in 10, 18 and 15 sites with maximum daily intensity of use of 0.65, 2.3 and 6.3 respectively (Fig. S5).

The final dataset comprised of a large number of variables including the vegetation (canopy, tree richness, DBH avg, total density, reg density (ha), grass, forbes, moss, ivy, litter and seedlings), soil properties (pH and OM), vertebrates (badger, bird, cat, dog, fox, hedgehog, lagomorph, micromammal, mustelid and squirrel encounter rates and detections in sound recorders), soil microarthropod fauna (24 groups), the diversity, composition and taxa relative abundance of bacteria and fungi (high through-put sequencing data, with various thousands of ASVs and hundreds of species). Multivariate ordination showed no clear clustering of soil samples based on deer presence or absence for soil microbes, either bacteria or fungi (Fig. S6). Similarly, the alpha-diversity indices showed non-significant impact of deer on bacterial and fungal Shannon diversity and species richness (Fig. S7). Also, PLFAs show no responses. Nevertheless, DESeq2 analysis identified 76 bacterial ASVs (corresponding to 38 genus-level taxa) and 82 fungal ASVs (corresponding to 53 genus-level taxa) which were significantly affected by deer presence or absence (Fig. 2).

**Fig. 2.**
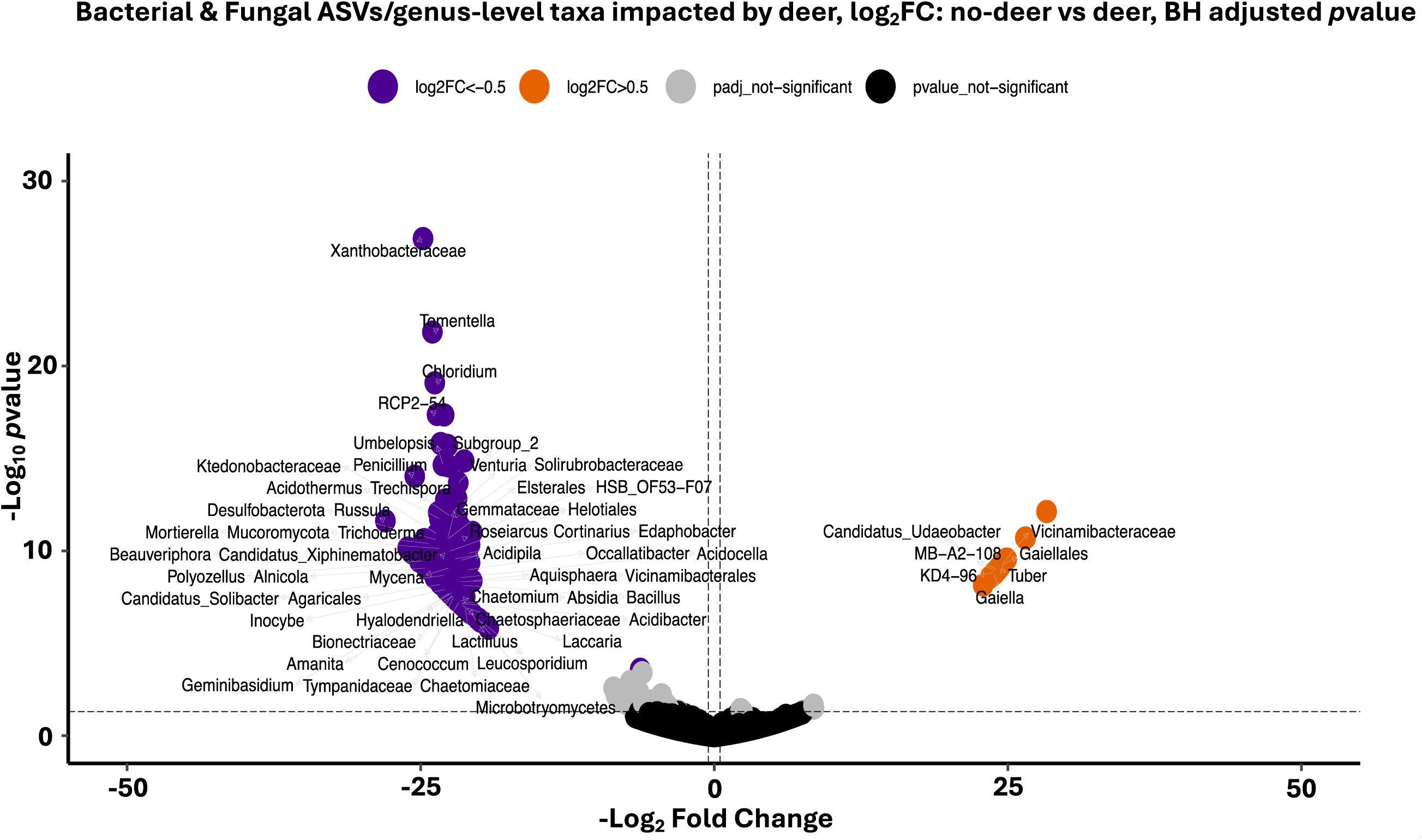
Volcano plot of the deer-dependent changes in the bacterial and fungal communities of soil identified by Deseq2 analysis. Genus name was used as label or, where Genus could not be identified, family or order names were used. Orange dots indicate genera dominant in no-deer, while purple in deer samples.

Principal Coordinate Analysis was then applied to each individual data. The first PCoA axis labelled as PC1 was used as a descriptor of the various groups of variables (bacteria, fungi, soil fauna, vertebrate fauna and vegetation). In each ordination, PC1 accounted for 22% of the variation in the vegetation data, 26% of the variation in vertebrates, 40% of the variation in soil microarthropods, and 23% and 35% of variations in selected bacterial and fungal communities’ data respectively (Fig. S8). Despite this amount of variation being around 25% and 40% of variance, it was in many cases well correlated to our metric of deer intensity of use at the site level as well as with other co-variables known to affect the various measured variables (for example pH vs bacteria). For example, redundancy analysis (RDA) returned two significant (*P* < 0.05) variables (pH and soil fauna) accounting for 10.0% (adjusted R^2^) of the variance in bacterial community composition (Fig. 3a). For fungal data, the significant variables were pH and vertebrates (*P* < 0.05), accounting for 7.8% of the total variation in fungal community composition (Fig. 3b). Soil fauna composition across sites were mainly affected by pH and bacteria (*P* < 0.01), which accounted for 14.3% of the total variation (Fig. 3c). The composition of vertebrates was significantly affected just by deer (Fig. 3d).

**Fig. 3.**
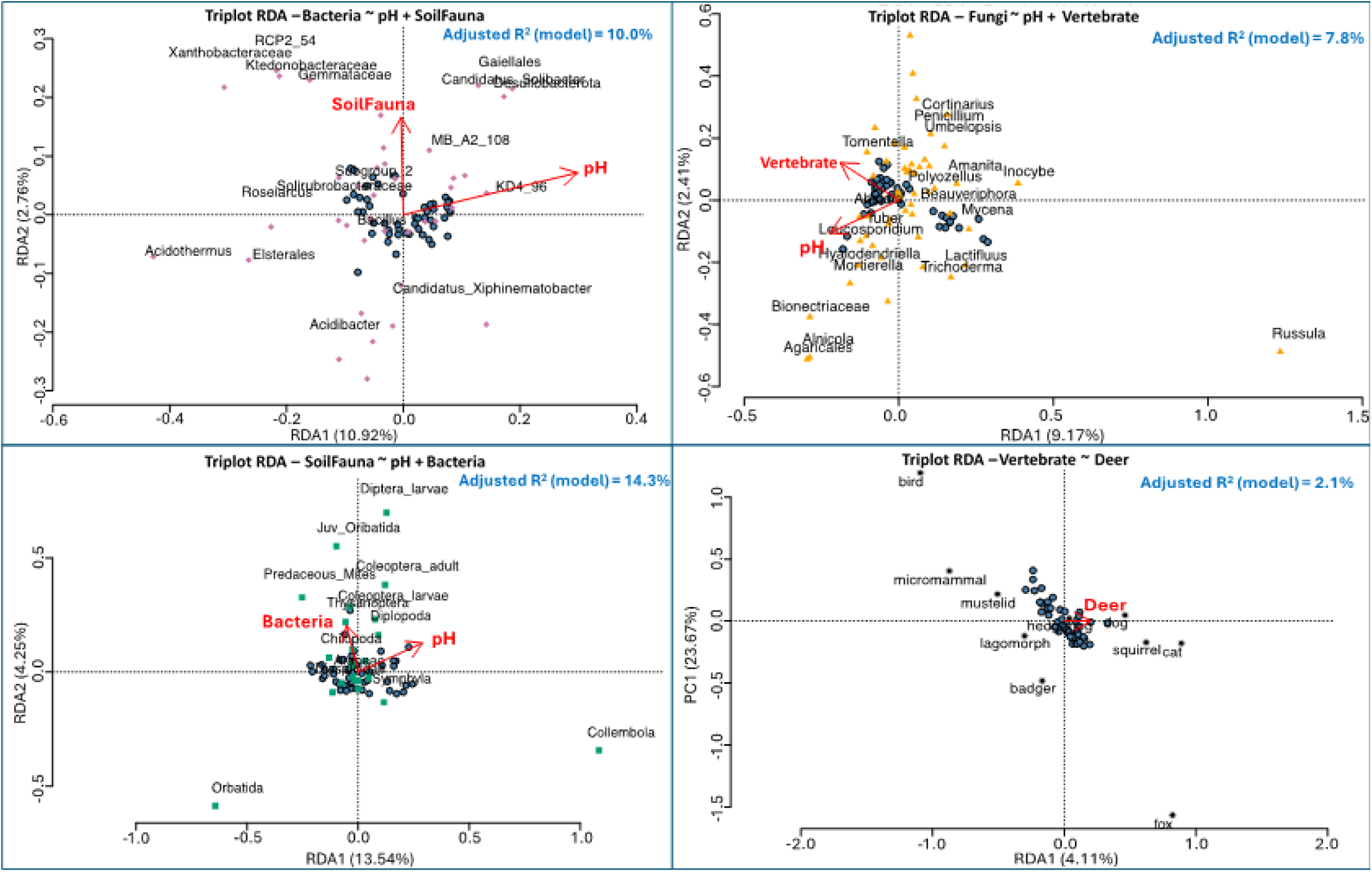
Redundancy analysis using forward selection method to model the effect of statistically significant variables on the selected communities of bacteria (a), fungi (b), soil fauna (c) and vertebrates (labelled as megafauna) (d). In each plot light blue dots are samples (sites). The significant explanatory variables are shown with red arrows, while the arrow’s length corresponds to the strength of the correlation . The % of the variance explained (adjusted R2) in species abundances of bacteria, fungi, soil fauna and megafauna across sites is shown.

SEM model 3 (with indirect effects only) was selected as the best model based on the goodness of fit determination (including non-significant chi-square and lowest AIC value of 287.239 (Table S2) as compared to the AIC values of model 2 (293.943) and model 1 (292.516) respectively (See also Figure S9 for model 1 and 2, and Table S1 for full list of models’ parameters). SEM further confirmed that varying deer densities directly affected the community composition of vertebrates (R^2^ = −0.32, *P* < 0.05), where the intensity of use by deer was positively associated with fauna such as cat, fox, squirrel, dog, while negatively associated with bird, micromammal, mustelid and lagomorph (Fig. 4). Deer intensity of use also covaried with vegetation through an increase in litter, moss quantity, while a decrease in forbes, ivy, reg density (ha), seedlings, DBH avg, tree richness, canopy cover, and grass (R^2^ = −0.20, *P* = 0.17, Fig. S8) . These effects cascaded indirectly to soil biota (soil fauna, bacteria and fungi) through changes in soil pH. Bacterial communities were strongly affected by pH (R^2^ = 0.62, *P* < 0.001), followed by soil fauna (R^2^ = −0.38, *P* < 0.001). Both pH and soil fauna were positively associated with the abundances of bacterial genera such as *Desulfobacterota* and *Gaiellales*, while negatively associated the abundances of *Acidothermus, Xanthobacteraceae, Elsterales, Roseiarcus, Ktedonobacteraceae, RCP2_54 and Gemmataceae* (Fig. 4). Both pH and vertebrate fauna equally effected the fungal communities (similar path coefficients; see also supporting information Table S1 for full list of model parameters) and negative R^2^ value of −0.33 (*P* < 0.05). A positive relationship of pH and vertebrate fauna with the abundances of fungal genus-level taxa such as *Agaricales, Alnicola*, and *Bionectriaceae*, while a negative relationship with the abundances of *Russula, Inocybe, Mycena, Lactifluus*, *Amanita, Trichoderma* and *Beauveriphora* were observed. In case of soil fauna, a strong positive effect of pH on Collembola community, while a strong negative effect on the communities Oribatida and predaceous mites was observed (Fig. 4).

**Fig 4.**
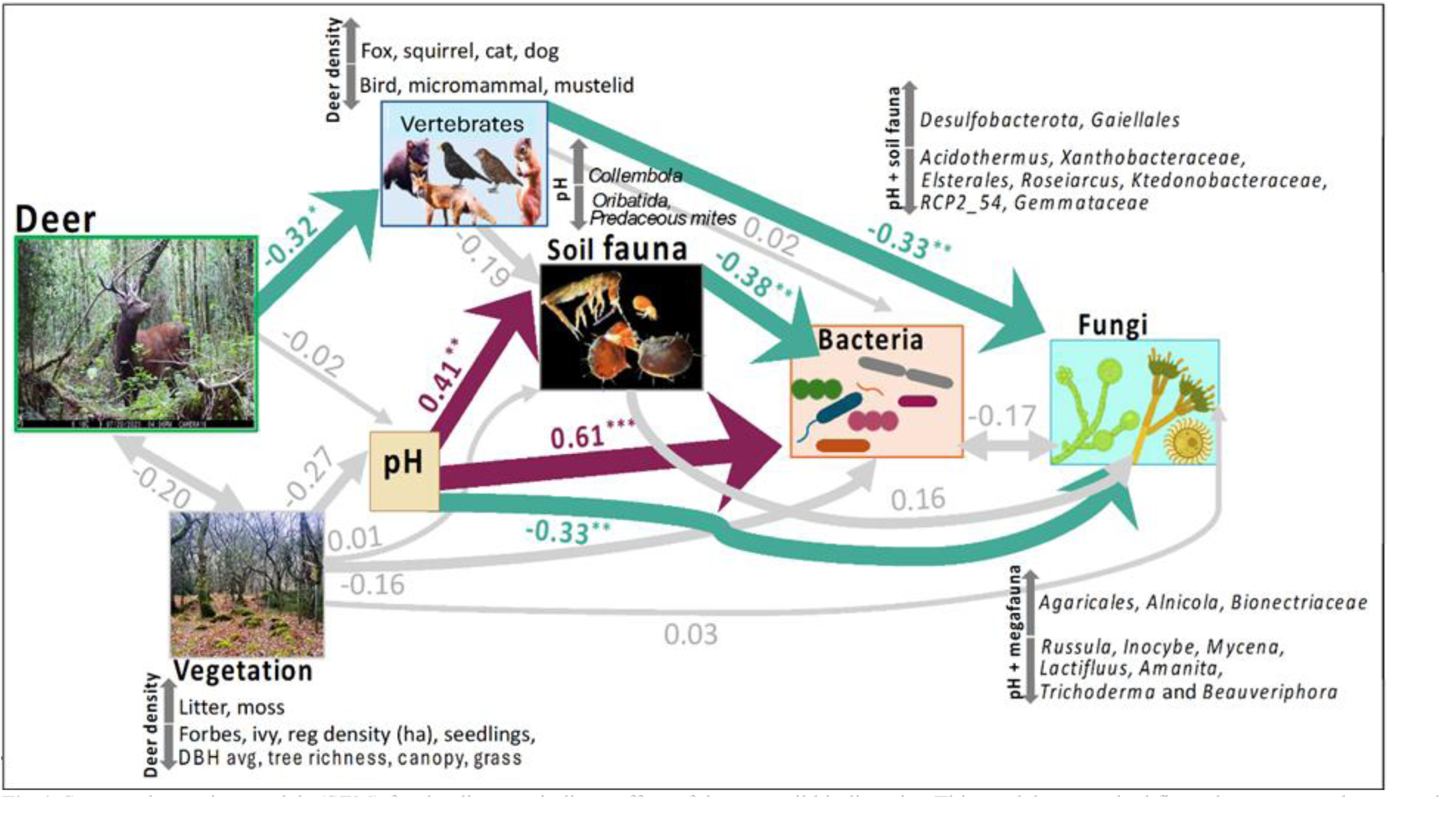
Structural equation models (SEM) for the direct vs indirect effect of deer on soil biodiversity. This model was ranked first when compared to two other model structures (Supporting Information): one with just direct effects, and one with just indirect effects. Arrow thickness corresponds to the value of standardised coefficients for each path (as shown). Paths shaded strong pink indicate significant positive associations, while, those shaded green indicate significant negative associations. Paths with non-significant *P*-value are shaded grey.

## 4. Discussion

Overall, our study showed clear empirical evidence that high deer site-level intensity of use indirectly affects soil biota, via changes in soil pH and vertebrate fauna, alongside covariation with vegetation structure. That is partially in contrast with some other studies documenting both direct and indirect effects of deer on the composition and diversity of soil fauna through habitat destruction by trampling resulting in reduced soil porosity [59], a reduction in litter quality by deer intensity of use leading to reduced soil fauna activity and abundance [60], or long-term deer grazing-induced alterations in the quality and quantity of litter, which can favour Collembola but reduce abundance of predacious mites [61]. An explanation of the dominance of indirect effects in our study is exactly that we considered also vertebrate fauna: in our case, higher deer intensity of use was associated with a decrease in the abundances of birds, micromammals, mustelids and lagomorphs and an increase in anthropogenic-related faunas. That change in vertebrate fauna cascaded, in terms of the correlations modelled with the SEM, on soil bacteria and fungi but not on soil microarthropods, which resulted mostly driven by changes in soil pH and vegetation, and with the latter well correlated to deer intensity of use. Regardless of whether the effects were direct or indirect, we, too, found that study sites with higher deer intensity of use were dominated by Collembola, while Oribatida and predaceous mites in the Mesostigmata did the opposite. We attribute this anticorrelation pattern to a deer indirect effect because our structural equation model selection offers empirical evidence the effect of the deer was linked to covariation between pH, vegetation and deer intensity of use.

Soil pH is a known determinant of soil fauna communities [62]. Feketeová and colleagues also highlighted the importance of soil pH as a controlling factor of forests colonization by macro decomposers such as exotic earthworms, with an essential role in litter layer decomposition [63], thus, indirectly affecting the communities of oribatid mites [64]. That suggests that the effect of pH triggered by deer intensity of use may itself operate indirectly on soil fauna through biota we did not measure, such as earthworms and enchytraeids. For example, collembolans feed on soil organic matter and associated soil microorganisms [65]. Changes in soil pH alter the community composition of soil microorganisms (something confirmed by our data as well) which can then indirectly affect the composition of soil collembolans [66, 67]. The collembola community can also directly modify the composition of soil bacterial and fungal communities through selective consumption [68]. Hågvar and Abrahamsen [69] also reported a positive association of collembolans with soil pH in forest sites. The water absorption ability of collembolans is also pH dependent, with low pH resulting in dehydration and death [70]. On the contrary, negative effects of alkaline soil pH on the abundance and diversity of oribatid mites were recently reported [64, 71]. Likewise, studies on predacious mites also reported their higher abundance in low pH soils [72, 73]. Overall, the data suggest that the main effect of deer on soil microarthropods was mostly indirect, through covariation with vegetation yet consistent with what expected in terms of the response of the three major groups of soil microarthropods to pH changes.

Instead, the indirect effect of deer on microbes was through the effects of deer on vertebrate fauna, as well as through the covariation with vegetation and pH. Soil microbial communities play an essential role in the carbon and nitrogen cycling [74] and are highly sensitive to environmental changes due to their close link to the vegetation and understory layers, nutrient gradients and soil inhabiting organisms [75–77]. Deer herbivory can directly affect the diversity and composition of plant species and indirectly affects soil properties and the associated soil microorganisms [78] which is similar to our data suggesting that the effects are indirect. Also, low soil pH may favour the development of fungi and a decrease in fungi to bacteria ratio [79, 80] but in our case that ratio, as measured by PLFAs, did not respond. Soil pH was also reported to be the major factor contributing to the variance in bacterial community composition across a range of studies [81–83]. We, indeed, found a high abundance of *Acidothermus, Xanthobacteraceae, Elsterales, Roseiarcus, Ktedonobacteraceae, RCP2_54,* and *Gemmataceae* in highly acidic soil, which were also those with higher deer intensity of use, while a high abundance of *Desulfobacterota* and *Gaiellales* in sites with high soil pH and in the absence of or lower intensity of use (Fig. 4 & Fig. S8). *Acidothermus* is an acidophilic bacterial genus, greatly impacted by soil pH [84], showing relatively high abundance under acidic soil condition [85, 86]. *Roseiarcus* is a symbiotic bacterium, known for its associations with fungi in the mycosphere to promote plant growth [87]. A high abundance of *Roseiarcus* was observed in the coniferous forest with very low soil pH [84, 88]. Similar to our findings, an association of the genera *Acidothermus, Roseiarcus, Ktedonobacteraceae, Elsterales* and *RCP2_54* with low soil pH values [89] and a negative association between *Xanthobacteraceae* abundance and soil pH was recently reported [90]. As our data are fully consistent with the literature, the implication of our model is that any variation in soil pH caused by deer either directly or indirectly through changes in vegetation density and composition (e.g., deer trampling), even if weak and only indirect, can affect microbial composition and so the functions expressed by them.

A similar effect was observed for soil fungal communities. We observed a higher abundance of ectomycorrhizal fungi (*Russula, Inocybe, Lactifluus, Amanita*), plant and animal pathogenic fungi (*Mycena* and *Trichoderma*) in the more acidic soil which had high deer intensity of use, while high abundances of saprotrophic, bryophyte and algal parasite such as *Alnicola, Agaricales* and *Bionectriaceae* with an increase in soil pH . Soil pH is known to be a strong predictor of the community composition of soil fungi (e.g. [91]) as it can affect the growth of ectomycorrhizal fungi, production of fruiting body and its distribution thus, effecting plant productivity and growth [92]. A pH range of 4-5 [93], and 5-6 [94] were found optimum and most suitable for the growth and root colonization of ectomycorrhizal fungi. However, pH range of 7-8 was most suitable for growth of saprotrophic fungal species [94]. Likewise, short-term increase in soil pH was found to favour opportunistic groups proliferation rather than ectomycorrhizal fungi [95]. Once again, the implication is that changes in soil pH linked to variation in deer intensity and deer effect on vegetation have a strong potential to affect the soil fungal community.

Deer directly interacts with fungi: mushrooms, the fungus fruiting body, contribute 5-15% to the diet of Cervidae because of their high digestibility and protein content [96, 97]. About 50 species of mushroom and lichen were identified in the Cervidae diet [98]. Few studies reported voluntarily selection and consumption of mushrooms by cervids when available [99, 100]. Certain plant pathogenic fungi and fungal endophytes present in plant tissues are also consumed by cervids [101]. In addition, deer may also influence composition of fungal communities through spores’ dispersal [102]. We found high abundances of ectomycorrhizal fungal genera (*Russula, Inocybe, Lactifluus* and *Amanita*) in high deer intensity of use sites, which might suggest their seasonal preference in sika deer diet as reported in other studies. *Russula* is one of the most diverse and largest basidiomycete genera, known for its wide geographical and ecological distribution and its ability to form ectomycorrhizal associations with a variety of plants [103, 104]. *Russula* is rich in protein content and was reported to be a preferred component in the diet of white-tailed deer (*Odocoileus virginianus*) [98]. *Russula, Amanita*, and *Inocybe* were among the few important fungal genera that were found in roe deer (*Capreolus capreolus*) diet [105]. In addition to the ectomycorrhizal fungi, our study also reported high abundances of plant and animal pathogenic fungi including *Mycena* and *Trichoderma* with high deer intensity of use. Similar to our study, Kadowaki, Honjo [106] reported high prevalence of animal and plant pathogenic fungi in the presence of deer resulting in high urine and faecal deposition as compared to fenced sites. Overall, our studies offer strong evidence that deer can have a major impact on fungi. Intriguingly, however, our statistical models suggest mainly an indirect effect, despite the potential for direct effects highlighted by previous studies. As the indirect effects was linked to both pH and other vertebrate fauna, which might arguably also directly affect fungi, future studies should more mechanistically tease apart the relative roles of indirect and direct effects.

Soil biota play a crucial role in degradation of organic matter and mineralization, symbiosis with plants as well as pathogens, which are all involved in decomposition of residual plant and animals resulting in biomass production that can be utilized by plants and soil organisms themselves [107–110]. Our data overall show evidence that in deer hotspot areas, the community composition of soil biota can vary in response to variation in vegetation structure and features such as soil pH, which are known key determinants [75, 78, 111]. The effect of deer intensity of use on other vertebrate fauna also seems to have a role, with that fauna becoming more anthropic in areas with high deer intensity of use – this could be the end result of reduced understory vegetation and increased accessibility of these sites by humans and human-related fauna, although this requires further investigation. That effect mimics changes typical of soil biota with relatively higher abundances of collembolan and opportunistic microbes.

While our study lacks the mechanistic resolution needed to resolve cause-effect relationships and functional implications of deer overabundance, the results we collected empirically demonstrate the potential for that overabundance to significantly alter soil biota community structure. Future studies will have to experimentally tease apart direct and indirect effect of deer populations on soil biota community structure and functions to assess the ecosystem level implications of altered aboveground-belowground linkages.

## Acknowledgements

We are thankful to all agencies and contributors that aided this project and allowed us to access the lands they manage. This project was funded by the Department of Agriculture, Food, and the Marine (DAFM) – grant (UCD grant R24354; 2023 DAFM Policy and Strategic Studies Research Call PSSRC). CB was jointly funded by DAFM and by Taighde Éireann – Research Ireland postgraduate scholarship. We thank Tony Quinn and Cian O’Connor for their invaluable and continued support throughout the project. We thank Barry Coad and Mary Clifford of Coillte for facilitating the selection and access to sampling stations. We thank Wesley Atkinson, Ann Fitzpatrick, Damian Clarke and Jenni Roche of the National Parks and Wildlife Service for their advice, input and help with selecting further sampling stations. We are thankful to Katherine Stafford, Mary Walsh, John Kavanagh, Cristin Deiseach, Brian Miller, Philip Hayden, Stephen Toth, William Bunbury, Bryanna Alton, Emmet Altidore, Richard Nairn, Lucy Tottenham, Zef Klinkenbergh and Ruth Hussey who so willingly provided us access to their private property to conduct this study across Wicklow and beyond. We thank Kilian Murphy for input during the initial brainstorming stage of the project. We thank everyone who joined in the fieldwork throughout the summer including Andrew Ryan, Beth Diamond, Fiachra Corcoran, Grace Nolan, Jane Faull, Mattie Purinton, Niamh Collins, and Yujing Zhou. Additional thanks to the School of Biology and Environmental Sciences, University College Dublin for their continued support. Finally, we thank the various foresters, farmers and enthusiastic individuals who showed an interest in the project and gave valuable feedback.

## Authors’ Contribution

CB and SC designed the experiment with contribution from TC, AFS and VMP. CB and AB conducted field work and processed soil samples. NA performed molecular analysis and bioinformatics to profile microbial communities. CB and NA performed statistical analysis with input from TC, VMP and SC. NA and CB drafted the manuscript with contribution from all authors.

## Notes

### Competing Interest Statement

The authors have declared no competing interest.

