## Supplemental Material for "Dominance of indirect effects of deer populations on soil biodiversity"

**Supporting Information**

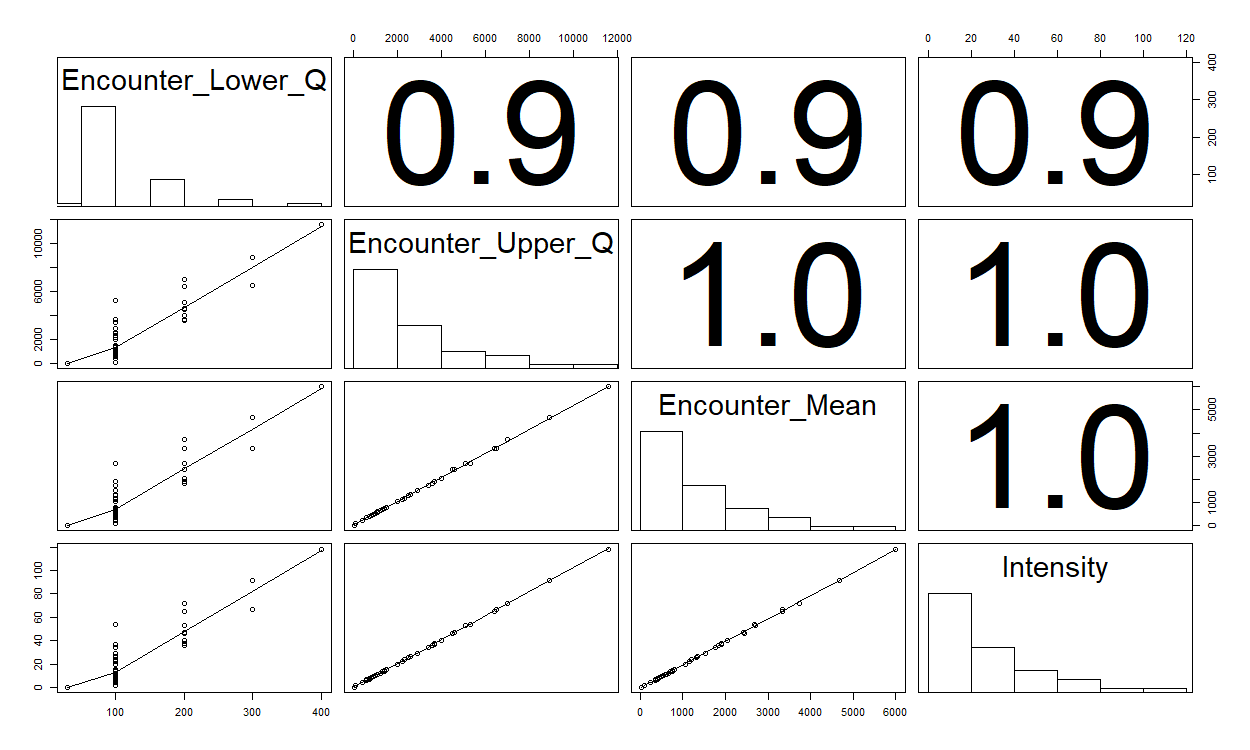

**Figure S1**: Correlation matrix showing Pearson Correlation Coefficients between deer encounter rate (lower quantile, upper quantile and mean) and intensity of use at camera trap sites.

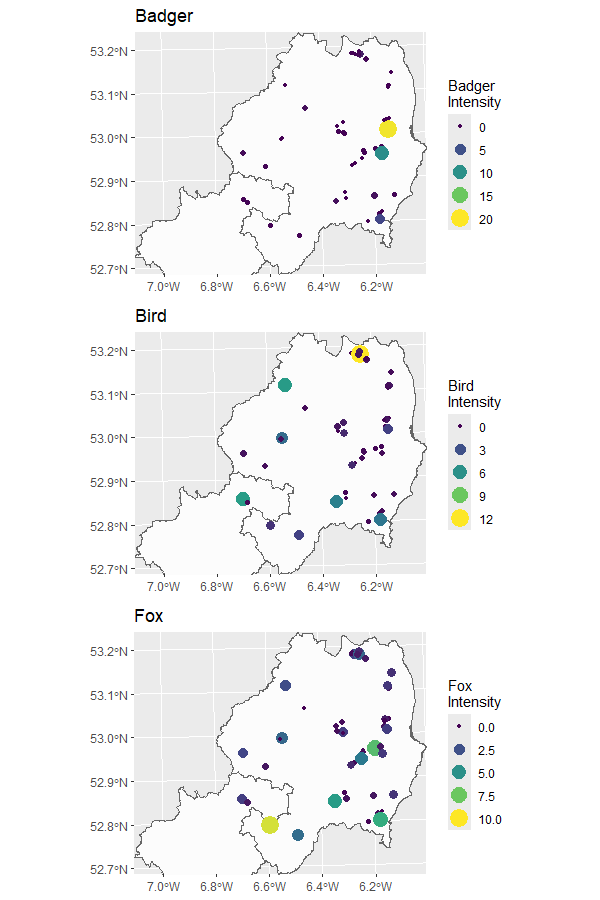

**Figure S2.** Spatial distribution of badger, bird and fox daily intensity of use within Wicklow and Carlow regions. Each point represents a sampling site with the size and colour highlighting the differences in daily intensity of use. Smaller and darker colours represent low daily intensity use and larger brighter colours represent higher daily intensity use (note the difference in scale for each species).

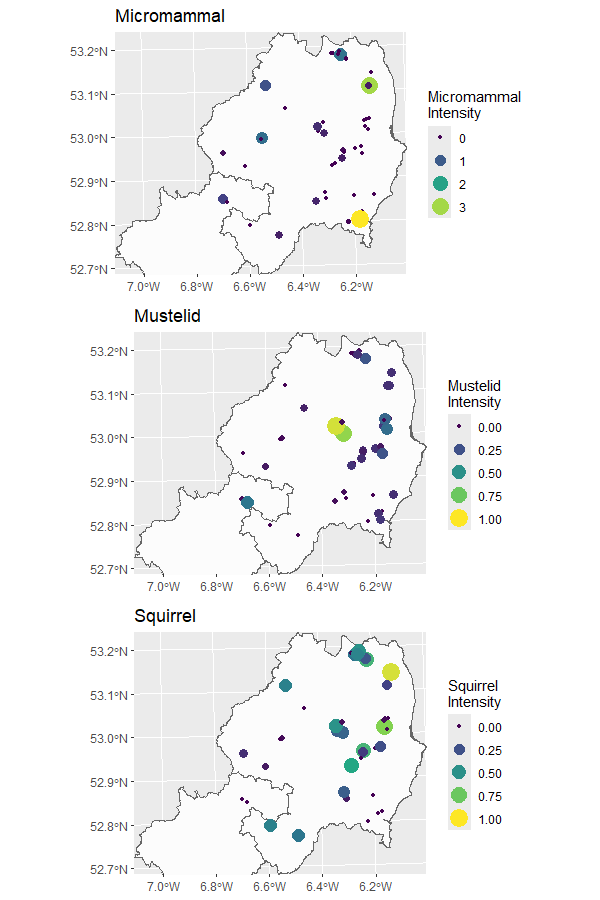

**Figure S3**. Spatial distribution of micromammal, mustelid and squirrel daily intensity of use within Wicklow and Carlow regions. Each point represents a sampling site with the size and colour highlighting the differences in daily intensity of use. Smaller and darker colours represent low daily intensity use and larger brighter colours represent higher daily intensity use (note the difference in scale for each species).

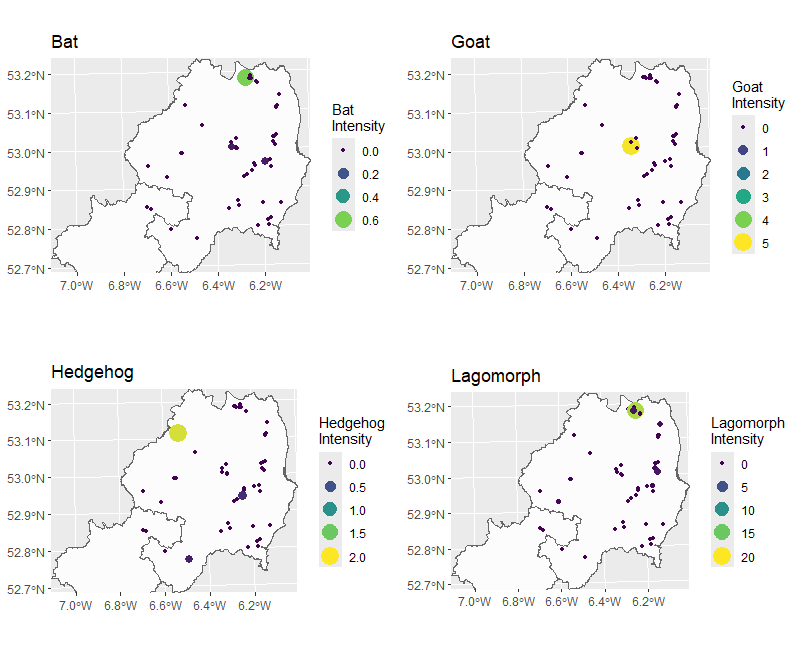

**Figure S4.** Spatial distribution of bat, goat, hedgehog and squirrel daily intensity of use within Wicklow and Carlow regions. Each point represents a sampling site with the size and colour highlighting the differences in daily intensity of use. Smaller and darker colours represent low daily intensity use and larger brighter colours represent higher daily intensity use (note the difference in scale for each species).

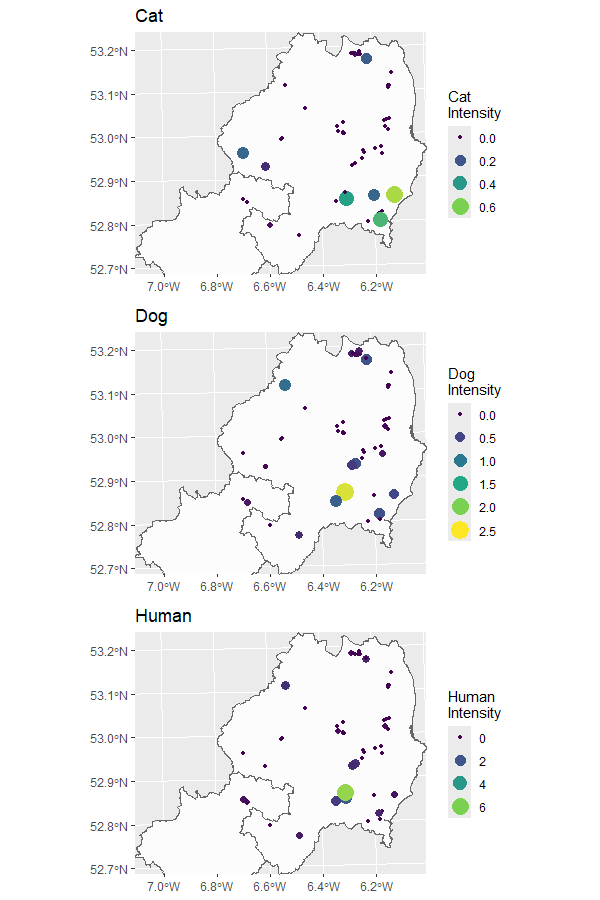

**Figure S5.** Spatial distribution of cat, dog, and human daily intensity of use within Wicklow and Carlow regions. Each point represents a sampling site with the size and colour highlighting the differences in daily intensity of use. Smaller and darker colours represent low daily intensity use and larger brighter colours represent higher daily intensity use (note the difference in scale for each species).

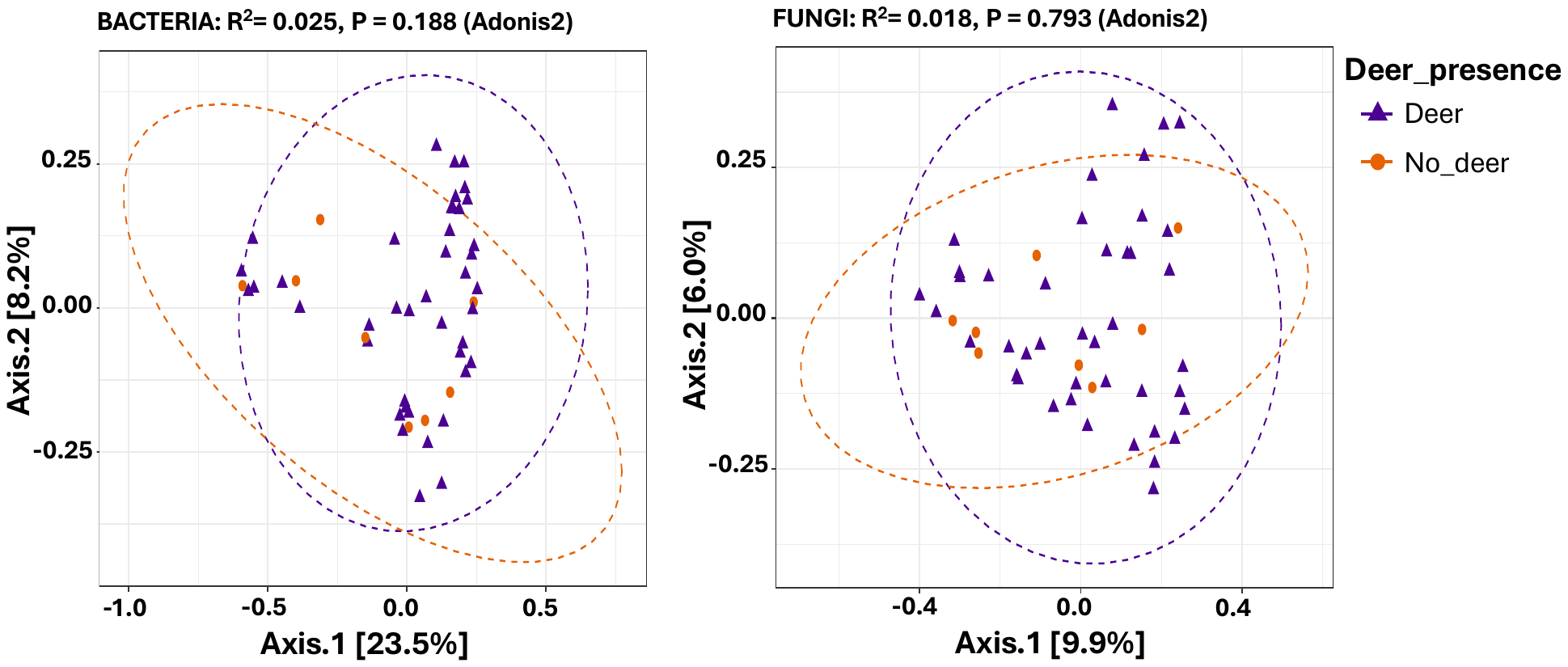

**Figure S6.** Beta-diversity analysis of soil samples collected from 50 sites across co. Wicklow. Beta-diversity was performed at ASVs level with Bray-Curtis as similarity matrix and Principal Coordinate Analysis (PCoA) plot as visualisation method.

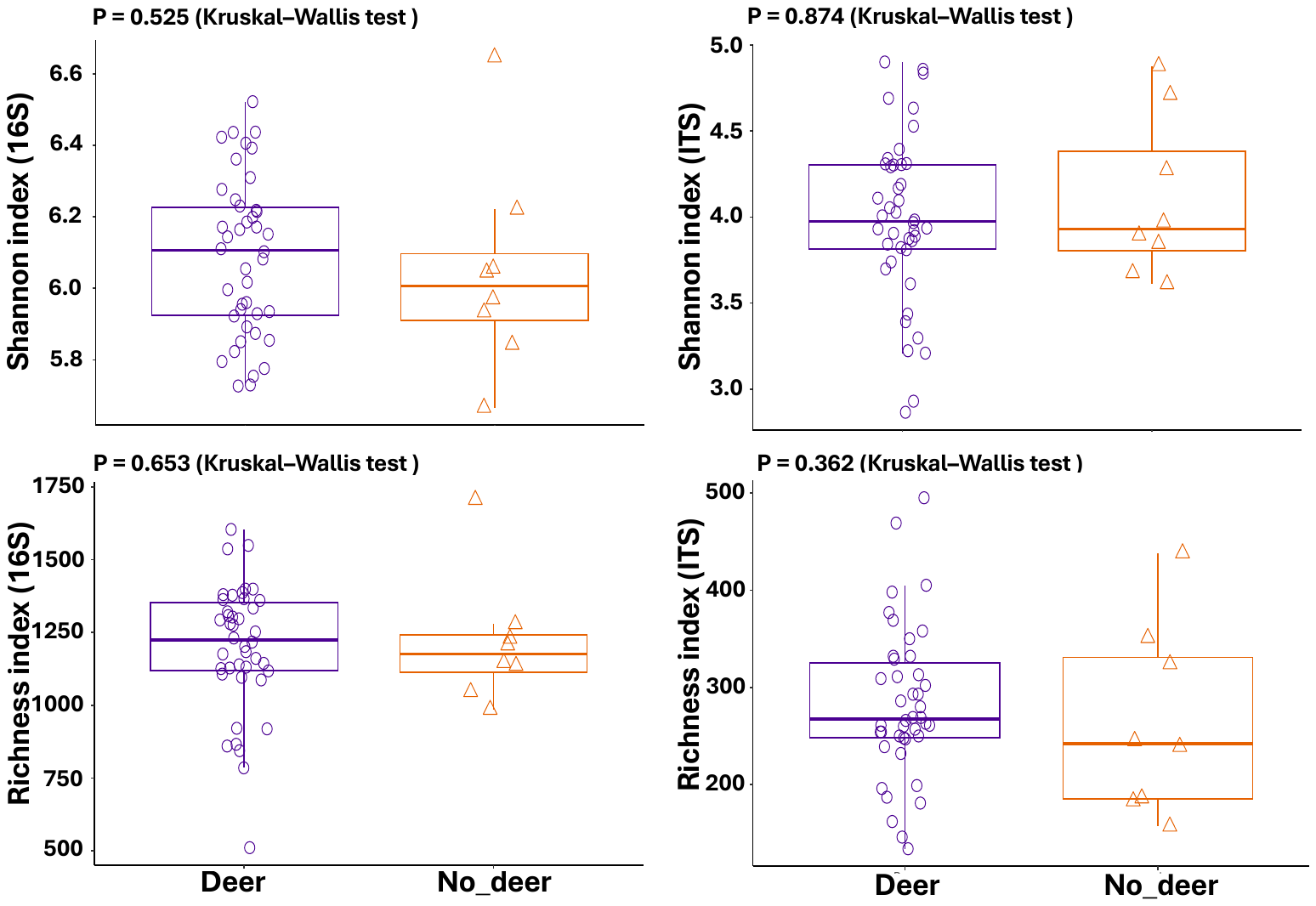

**Figure S7.** Alpha-diversity analysis of soil samples collected from 50 sites across co. Wicklow. Alpha-diversity was performed on standardised and rarefied ASVs count data. P values are based on Kruskal–Wallis test.

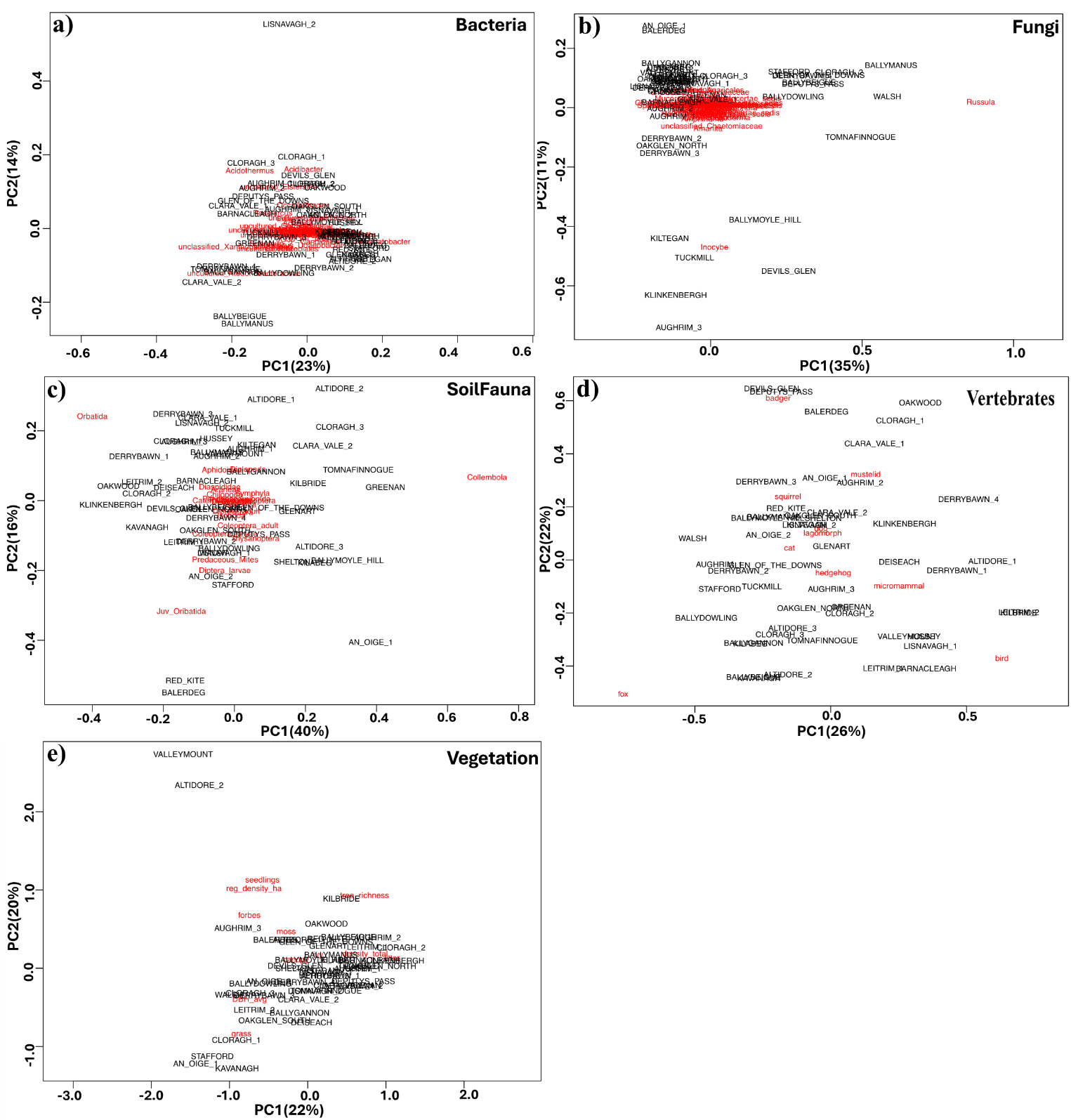

**Figure S8.** Principal Component Analysis (PCA) plots to find associations of different sites (samples in black) with particular variable (bacteria, fungi, soil fauna, vertebrates and vegetation).

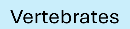

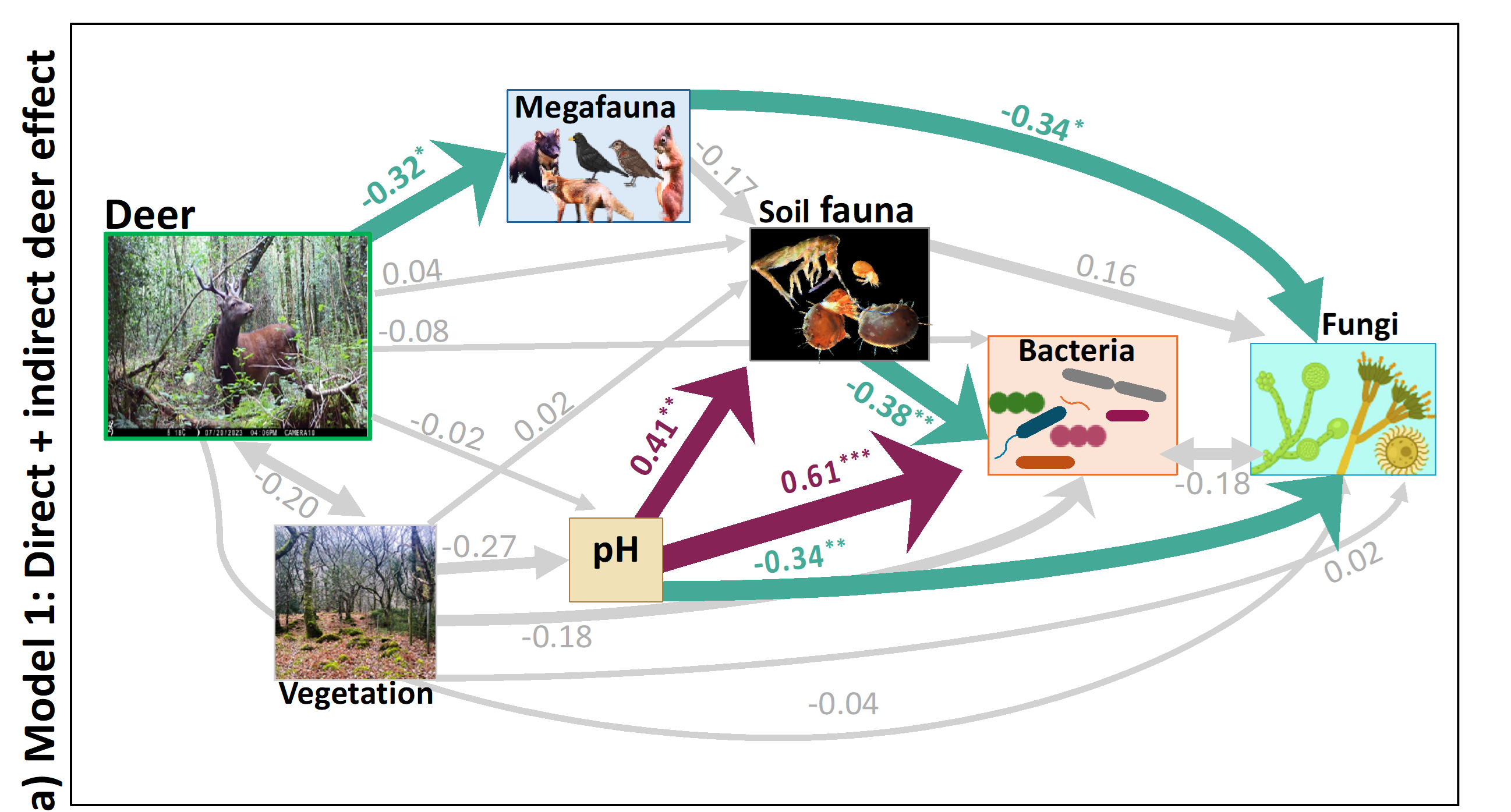

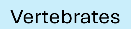

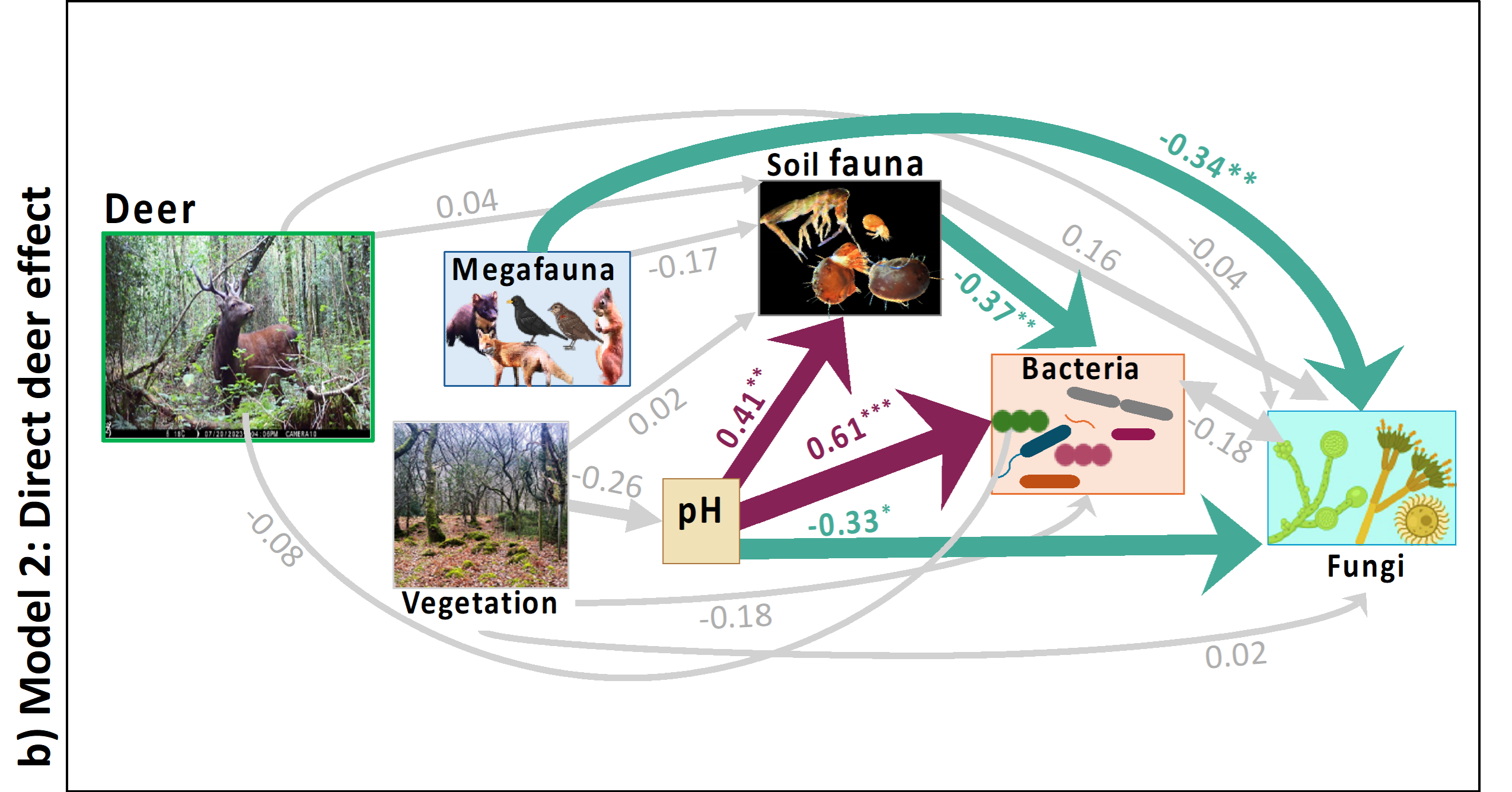

**Figure S9.** SEM, model 1 (a) and model 2 (b). Model 3 is illustrated in Figure 4 of the main manuscript.

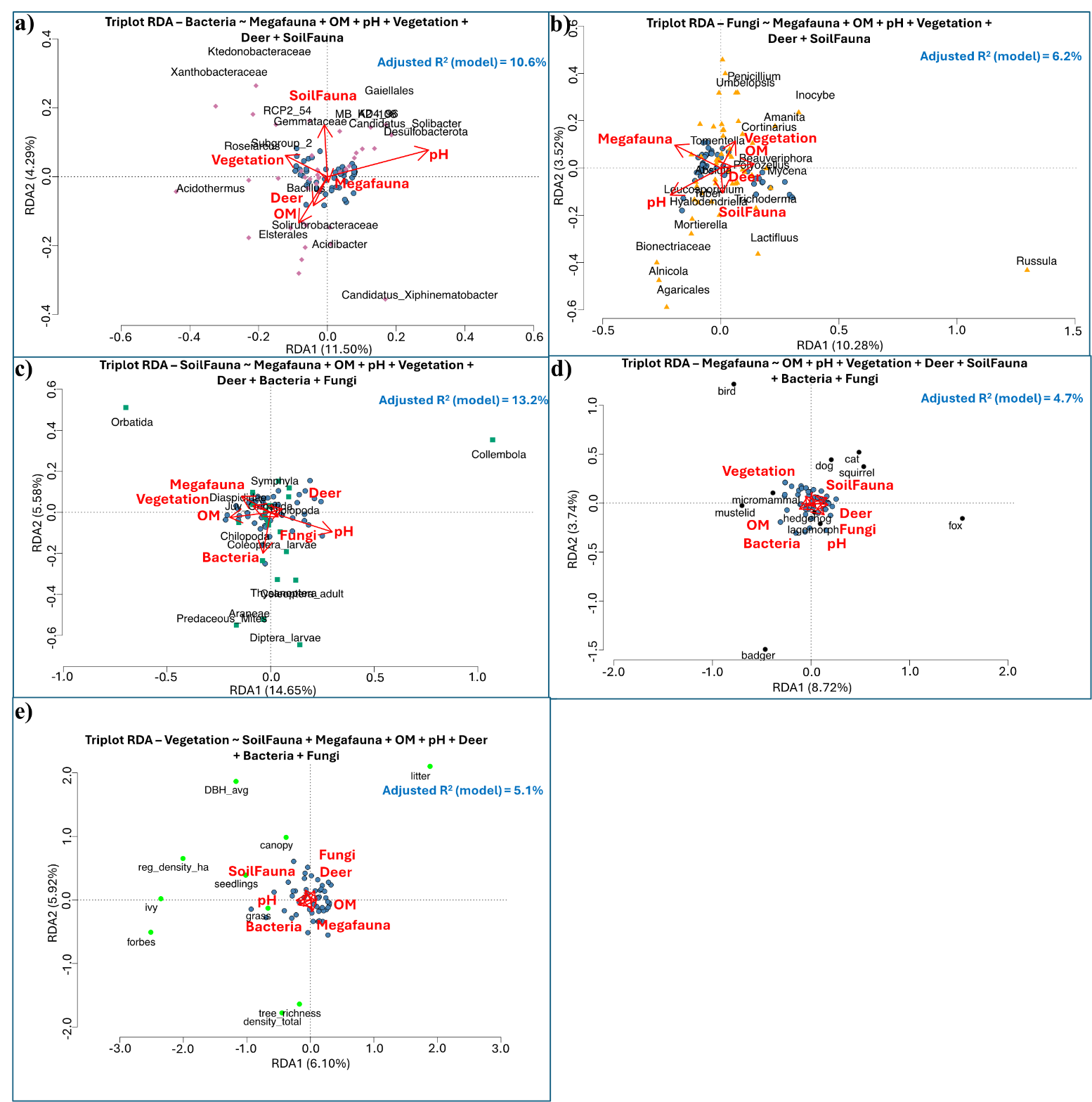

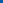

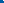

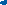

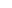

**Vertebrate**

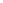

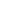

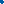

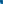

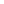

**Vertebrate**

**Triplot RDA – Vegetation ~ SoilFauna + Vertebrate + OM + pH + Deer + Bacteria + Fungi**

**Triplot RDA – Vertebrate ~ OM + pH + Vegetation + Deer + SoilFauna + Bacteria + Fungi**

**Triplot RDA – SoilFauna ~ Vertebrate + OM + pH + Vegetation + Deer + Bacteria + Fungi**

**Vertebrate**

**Triplot RDA – Fungi ~ Vertebrate + OM + pH + Vegetation + Deer + SoilFauna**

**Vertebrate**

**Triplot RDA – Bacteria ~ Vertebrate + OM + pH + Vegetation + Deer + SoilFauna**

**Figure S10.** Redundancy analysis (RDA) to model the effect of each set of variables (both significant and non-significant) on the selected communities of bacteria **(a)**, fungi **(b)**, soil fauna **(c)**, vertebrates **(d)** and vegetation **(e)**. Light blue dots are samples (sites). The explanatory variables and the **%** of the variance explained (adjusted R^2^) in species abundances of bacteria, fungi, soil fauna, megafauna and vegetation across sites is shown.

**Table S1.** The full output of the Structural Equation Models (SEMs) graphically presented in Figure 4 is provided below:

**Model 1:**

| Regressions: | |  |  |  |  |  |
| --- | --- | --- | --- | --- | --- | --- |
|  | Estimate | Std.Err | z-value | P(>\|z\|) | Std.lv | Std.all |
| pH ~ |  |  |  |  |  |  |
| Vegetation | -0.386 | 0.202 | -1.906 | 0.057 | -0.386 | -0.265 |
| Deer | -0.023 | 0.139 | -0.169 | 0.866 | -0.023 | -0.023 |
| Vertebrates ~ | |  |  |  |  |  |
| Deer | -0.096 | 0.04 | -2.385 | 0.017 | -0.096 | -0.32 |
| Fungi ~ |  |  |  |  |  |  |
| Vertebrates | -0.253 | 0.099 | -2.559 | 0.01 | -0.253 | -0.342 |
| pH | -0.075 | 0.031 | -2.381 | 0.017 | -0.075 | -0.337 |
| Vegetation | 0.007 | 0.042 | 0.163 | 0.871 | 0.007 | 0.021 |
| Deer | -0.009 | 0.03 | -0.304 | 0.761 | -0.009 | -0.041 |
| SoilFauna | 0.185 | 0.164 | 1.131 | 0.258 | 0.185 | 0.158 |
| Bacteria ~ |  |  |  |  |  |  |
| Vertebrates | 0.001 | 0.049 | 0.011 | 0.991 | 0.001 | 0.001 |
| pH | 0.076 | 0.015 | 4.936 | 0 | 0.076 | 0.612 |
| Vegetation | -0.032 | 0.021 | -1.542 | 0.123 | -0.032 | -0.178 |
| Deer | -0.01 | 0.015 | -0.669 | 0.504 | -0.01 | -0.079 |
| SoilFauna | -0.248 | 0.081 | -3.072 | 0.002 | -0.248 | -0.376 |
| SoilFauna ~ |  |  |  |  |  |  |
| Vertebrates | -0.109 | 0.084 | -1.299 | 0.194 | -0.109 | -0.173 |
| pH | 0.078 | 0.025 | 3.16 | 0.002 | 0.078 | 0.413 |
| Vegetation | 0.005 | 0.037 | 0.13 | 0.896 | 0.005 | 0.017 |
| Deer | 0.008 | 0.026 | 0.3 | 0.764 | 0.008 | 0.041 |
| Covariances: | |  |  |  |  |  |
|  | Estimate | Std.Err | z-value | P(>\|z\|) | Std.lv | Std.all |
| .Fungi ~~ |  |  |  |  |  |  |
| .Bacteria | -0.003 | 0.003 | -1.225 | 0.221 | -0.003 | -0.176 |
| Vegetation ~~ | |  |  |  |  |  |
| Deer | -0.134 | 0.097 | -1.377 | 0.168 | -0.134 | -0.199 |
| Variances: |  |  |  |  |  |  |
|  | Estimate | Std.Err | z-value | P(>\|z\|) | Std.lv | Std.all |
| .pH | 0.913 | 0.183 | 5 | 0 | 0.913 | 0.931 |
| . Vertebrates | 0.079 | 0.016 | 5 | 0 | 0.079 | 0.898 |
| .Fungi | 0.037 | 0.007 | 5 | 0 | 0.037 | 0.775 |
| .Bacteria | 0.009 | 0.002 | 5 | 0 | 0.009 | 0.596 |
| .SoilFauna | 0.028 | 0.006 | 5 | 0 | 0.028 | 0.795 |
| Vegetation | 0.464 | 0.093 | 5 | 0 | 0.464 | 1 |
| Deer | 0.98 | 0.196 | 5 | 0 | 0.98 | 1 |
| R-Square: |  |  |  |  |  |  |
|  | Estimate |  |  |  |  |  |
| pH | 0.069 |  |  |  |  |  |
| Vertebrates | 0.102 |  |  |  |  |  |
| Fungi | 0.225 |  |  |  |  |  |
| Bacteria | 0.404 |  |  |  |  |  |
| SoilFauna | 0.205 |  |  |  |  |  |

**Model 2:**

| Regressions: | |  |  |  |  |  |
| --- | --- | --- | --- | --- | --- | --- |
|  | Estimate | Std.Err | z-value | P(>\|z\|) | Std.lv | Std.all |
| pH ~ |  |  |  |  |  |  |
| Vegetation | -0.379 | 0.198 | -1.91 | 0.056 | -0.379 | -0.261 |
| Fungi ~ |  |  |  |  |  |  |
| Vertebrates | -0.253 | 0.094 | -2.696 | 0.007 | -0.253 | -0.34 |
| pH | -0.075 | 0.031 | -2.382 | 0.017 | -0.075 | -0.335 |
| Vegetation | 0.007 | 0.042 | 0.166 | 0.868 | 0.007 | 0.021 |
| Deer | -0.009 | 0.028 | -0.326 | 0.744 | -0.009 | -0.04 |
| SoilFauna | 0.185 | 0.164 | 1.131 | 0.258 | 0.185 | 0.157 |
| Bacteria ~ |  |  |  |  |  |  |
| Vertebrates | 0.001 | 0.046 | 0.012 | 0.991 | 0.001 | 0.001 |
| pH | 0.076 | 0.015 | 4.937 | 0 | 0.076 | 0.609 |
| Vegetation | -0.032 | 0.02 | -1.573 | 0.116 | -0.032 | -0.177 |
| Deer | -0.01 | 0.014 | -0.719 | 0.472 | -0.01 | -0.078 |
| SoilFauna | -0.248 | 0.081 | -3.072 | 0.002 | -0.248 | -0.373 |
| SoilFauna ~ |  |  |  |  |  |  |
| Vertebrates | -0.109 | 0.08 | -1.371 | 0.17 | -0.109 | -0.173 |
| pH | 0.078 | 0.025 | 3.161 | 0.002 | 0.078 | 0.414 |
| Vegetation | 0.005 | 0.036 | 0.133 | 0.894 | 0.005 | 0.017 |
| Deer | 0.008 | 0.024 | 0.323 | 0.747 | 0.008 | 0.041 |
| Covariances: | |  |  |  |  |  |
|  | Estimate | Std.Err | z-value | P(>\|z\|) | Std.lv | Std.all |
| .Fungi ~~ |  |  |  |  |  |  |
| .Bacteria | -0.003 | 0.003 | -1.225 | 0.221 | -0.003 | -0.176 |
| Vegetation ~~ | |  |  |  |  |  |
| Deer | 0 |  |  |  | 0 | 0 |
| Vertebrates ~~ | |  |  |  |  |  |
| Deer | 0 |  |  |  | 0 | 0 |
| .pH ~~ |  |  |  |  |  |  |
| Deer | 0 |  |  |  | 0 | 0 |
| Variances: |  |  |  |  |  |  |
|  | Estimate | Std.Err | z-value | P(>\|z\|) | Std.lv | Std.all |
| .pH | 0.913 | 0.183 | 5 | 0 | 0.913 | 0.932 |
| .Fungi | 0.037 | 0.007 | 5 | 0 | 0.037 | 0.767 |
| .Bacteria | 0.009 | 0.002 | 5 | 0 | 0.009 | 0.591 |
| .SoilFauna | 0.028 | 0.006 | 5 | 0 | 0.028 | 0.8 |
| Vegetation | 0.464 | 0.093 | 5 | 0 | 0.464 | 1 |
| Vertebrates | 0.088 | 0.018 | 5 | 0 | 0.088 | 1 |
| Deer | 0.98 | 0.196 | 5 | 0 | 0.98 | 1 |
| R-Square: |  |  |  |  |  |  |
|  | Estimate |  |  |  |  |  |
| pH | 0.068 |  |  |  |  |  |
| Fungi | 0.233 |  |  |  |  |  |
| Bacteria | 0.409 |  |  |  |  |  |
| SoilFauna | 0.2 |  |  |  |  |  |

**Model 3:**

| Regressions: | |  |  |  |  |  |
| --- | --- | --- | --- | --- | --- | --- |
|  | Estimate | Std.Err | z-value | P(>\|z\|) | Std.lv | Std.all |
| pH ~ |  |  |  |  |  |  |
| Vegetation | -0.386 | 0.202 | -1.906 | 0.057 | -0.386 | -0.265 |
| Deer | -0.023 | 0.139 | -0.169 | 0.866 | -0.023 | -0.023 |
| Vertebrates ~ | |  |  |  |  |  |
| Deer | -0.096 | 0.04 | -2.385 | 0.017 | -0.096 | -0.32 |
| Fungi ~ |  |  |  |  |  |  |
| Vertebrates | -0.244 | 0.094 | -2.589 | 0.01 | -0.244 | -0.33 |
| pH | -0.074 | 0.031 | -2.366 | 0.018 | -0.074 | -0.334 |
| Vegetation | 0.009 | 0.042 | 0.216 | 0.829 | 0.009 | 0.028 |
| SoilFauna | 0.183 | 0.164 | 1.118 | 0.264 | 0.183 | 0.156 |
| Bacteria ~ |  |  |  |  |  |  |
| Vertebrates | 0.01 | 0.047 | 0.214 | 0.831 | 0.01 | 0.024 |
| pH | 0.077 | 0.016 | 4.951 | 0 | 0.077 | 0.615 |
| Vegetation | -0.03 | 0.021 | -1.451 | 0.147 | -0.03 | -0.165 |
| SoilFauna | -0.25 | 0.081 | -3.089 | 0.002 | -0.25 | -0.379 |
| SoilFauna ~ |  |  |  |  |  |  |
| Vertebrates | -0.117 | 0.08 | -1.463 | 0.144 | -0.117 | -0.185 |
| pH | 0.078 | 0.025 | 3.147 | 0.002 | 0.078 | 0.412 |
| Vegetation | 0.003 | 0.036 | 0.082 | 0.934 | 0.003 | 0.011 |
| Covariances: | |  |  |  |  |  |
|  | Estimate | Std.Err | z-value | P(>\|z\|) | Std.lv | Std.all |
| .Fungi ~~ |  |  |  |  |  |  |
| .Bacteria | -0.003 | 0.003 | -1.191 | 0.234 | -0.003 | -0.171 |
| Vegetation ~~ | |  |  |  |  |  |
| Deer | -0.134 | 0.097 | -1.377 | 0.168 | -0.134 | -0.199 |
| Variances: |  |  |  |  |  |  |
|  | Estimate | Std.Err | z-value | P(>\|z\|) | Std.lv | Std.all |
| .pH | 0.913 | 0.183 | 5 | 0 | 0.913 | 0.931 |
| . Vertebrates | 0.079 | 0.016 | 5 | 0 | 0.079 | 0.898 |
| .Fungi | 0.037 | 0.007 | 5 | 0 | 0.037 | 0.777 |
| .Bacteria | 0.009 | 0.002 | 5 | 0 | 0.009 | 0.6 |
| .SoilFauna | 0.028 | 0.006 | 5 | 0 | 0.028 | 0.797 |
| Vegetation | 0.464 | 0.093 | 5 | 0 | 0.464 | 1 |
| Deer | 0.98 | 0.196 | 5 | 0 | 0.98 | 1 |
| R-Square: |  |  |  |  |  |  |
|  | Estimate |  |  |  |  |  |
| pH | 0.069 |  |  |  |  |  |
| Vertebrates | 0.102 |  |  |  |  |  |
| Fungi | 0.223 |  |  |  |  |  |
| Bacteria | 0.4 |  |  |  |  |  |
| SoilFauna | 0.203 |  |  |  |  |  |

**Table S2.** Parameters for goodness of fit determination of the three different SEM models.

|  | **Chisq** | **P-value (Chi-square)** | **CFI** | **RMSEA** | **AIC** |
| --- | --- | --- | --- | --- | --- |
| **Model 1** | 0.543 | 0.762 | 1.000 | 0.000 | 292.516 |
| **Model 2** | 7.971 | 0.158 | 0.929 | 0.109 | 293.943 |
| **Model 3** | 1.267 | 0.938 | 1.000 | 0.000 | 287.239 |
